# Directional variation in evolution: consequences for the fitness landscape metaphor

**DOI:** 10.1101/015529

**Authors:** Philip Gerlee

**Affiliations:** Mathematical Sciences, Chalmers University of Technology and Göteborg University

**Keywords:** directional variation, fitness landscape, configuration space

## Abstract

The concept of a fitness landscape, which describes the relation between genotype (or phenotype) and fitness as a surface on which the population climbs uphill towards local peaks, is a central conceptual tool in evolutionary biology. Inherent in this metaphor is the assumption that the distance between any two points can be defined in the same way as for Euclidean space. However, many of the processes which generate genetic variation, such as gene duplication and lateral gene transfer, are not symmetric by nature, but occur more readily in one direction than the other. This asymmetry is also found in phenotypes, for reasons associated with the genotype-phenotype map and developmental constraints. This article provides an overview of of these processes and suggest how existing methods can be used for incorporating these asymmetric processes when visualising fitness landscapes.

## 1. Introduction

The notion of a fitness landscape, or adaptive landscape†, was introduced some 80 years ago by Sewall Wright (1932), and has since become a central conceptual tool in evolutionary thinking. The metaphor relies on the fact that to each genotype (or phenotype) one can assign a fitness, which determines the reproductive success of the respective organism, and that the population through variation and selection performs a hill-climbing movement up local gradients in the landscape defined by the mapping from genotype to fitness. This might sound straight forward, but hidden in this statement is an assumption about distance or “nearness” between different genotypes (or phenotypes). If this does not hold, we are unable to define an ordinary notion of distance between different points in the landscape and the whole metaphor of a fitness “landscape” fails. In the following I will argue that in many instances this is actually the case, and that this has given an erroneous view of evolution as a simple optimisation process in which population constantly are driven up to peaks in the fitness landscape.

Firstly, we will review the historical development of the idea since its conception, and then describe the points of critique it has meet. We will then discuss the notion of distance or metric assumed in the metaphor, and how it relates to the genetic operators which generate variation among genotypes and consequently phenotypes. In section 4 we deal with the problems which arise in genotype-based fitness landscapes, and show that they, for a variety of reasons, such as unequal cross-over and lateral gene transfer, fail to capture the notion of a metric. We then move on to show how the notion of a metric fails in the case of phenotypic (trait-based) fitness landscapes, via for example developmental constrains and redundancy in the genotype-phenotype map. In the final section of the paper we introduce a new concept, the ‘dynamical fitness landscape’, which resolves the above difficulties and provides a more balanced view of evolution on fitness landscapes.

## 2. Different flavours of fitness landscapes

The original fitness landscape introduced by Sewall Wright was in part inspired by the realisation that in a population of diploid individuals only a very small fraction of all the possible genotypes are ever present in the population at one instant, or in other words, the population only inhabits a small fraction of the genotype-space. By assigning a fitness to each genotype, we can imagine the population moving as a cloud of points in a landscape, consisting of valleys and peaks, where the height at a given point is determined by the fitness of the genotype representing it. The variation produced in the population together with the force of natural selection would then shift the population up the steepest gradient in a hill-climbing fashion.

The problem that Wright saw with this analogy was that the population would inevitably get stuck at local fitness peaks, and to solve this problem he suggested a “shifting-balance theory”, in which the species would be divided into subspecies, that would together search the fitness landscape in a more efficient manner. More recent developments have however suggested that the problem of the population getting stuck is exaggerated or even non-existent, and stems from the comparison with a 3-dimensional landscape. Considering the interdependence of different genes, one probably needs to take a large number of genes (on the order of hundreds) into account for an accurate view of a fitness landscape, and the topology of such high dimensional and discrete spaces is quite different from our intuitions of 3-dimensional Euclidean space, possibly rugged and with ridges connecting peaks in the landscape (Kauffman & Levin, 1987).

The fitness landscape employed by Wright was of a genotypic character, where two genotypes would lie next to each other if they were accessible by a change in a single allele. The precise structure of the genotype space envisaged by Wright was never specified, and only in a later paper did he say that the: “genotypes are packed side by side in such a way that each is surrounded by genotypes that differ by only one gene replacement” (Wright, 1988). In later developments of the genotypic fitness landscape each axis represented one locus with as many values as possible alleles, giving a landscape with as many dimensions as the number loci considered.

The theory of genetic fitness landscapes has seen major developments in the last decades with the work of Kauffman & Levin (1987) who investigated the structure of the simplified version of fitness landscapes, namely uncorrelated rugged landscapes. The importance of non-viable genotypes or holes in the fitness landscape was investigated by Gavrilets (1997), who proposed it as a means of speciation. Finally, we would like to mention the work of Fontana & Schuster (1998) who brought the importance of the mapping from genotype to phenotype into view. They considered the evolution of RNA sequences which fold into secondary structures, and showed that due to the redundancy in the GP-map evolution occurs on what is termed “neutral networks” and that transitions in structure and thus fitness are sudden.

The genotypic fitness landscapes are however only one variety of fitness landscapes and other entities have been used for defining the dimensions in fitness landscape. A more general view of a fitness landscape is to define it as a space *X*, usually termed a configuration space, together with a function *f*: *X* → ℝ, which to each member (configuration) *g* ∈ *X* assigns a fitness *f*(*g*). For example, one alternative to using genotypes is to use population-level properties such as gene frequencies as configurations, however this turns out to be misleading since the population as a whole does not necessarily maximise its average fitness (Moran, 1964).

Another approach, pioneered by Simpson (1944) and followed by many others (Niklas, 1994; McGhee, 1999; Marshall, 2006), has been to take individual-level properties or phenotypic traits, such as branch length in the case of plant evolution (Niklas, 1994), as the axes defining the landscape. As with genotypic landscapes the dimensionality is given by the number of traits chosen, however the ordering along each axis is in this case unclear. A natural solution is to simply order the different version of the trait according to some metric of the trait itself, e.g. ordering the different possible branch morphologies according to their length. This however assumes something about the genetics and developmental process which gives rise to the trait, namely that small genetic changes give rise to small changes in phenotypes. Going from a short to a long branch should then in principle be possible by consecutive genetics mutations which alter the trait in a continuous manner. If this is not the case, then the movement of populations on the surface would be discontinuous and restrained to trajectories defined by genetic or developmental constraints, and the metaphor of a fitness landscape would hardly be a helpful for understanding the evolutionary process. This is one of my points of contention, and will be discussed in more detail in section 5b.

Yet another utilisation of the landscape metaphor involves the evolution of enzymes (Poelwijk *et al*., 2007). Here the axes represent the base pairs which code for the amino acids the protein consists of, and the height of the landscape corresponds to the enzymatic activity of the protein, or the actual reproduction rate of the organism which carries the enzyme-coding gene in question. With the biochemical techniques available today one has been able to investigate the structure of such landscapes, and the findings show that they exhibit complex structural features such as vallyes and plateaus. Another interesting finding is that the structure of the landscape depends on what scale the fitness is defined at. For example it was found that the enzymatic activity of the enzyme IMDH is single-peaked, but when viewed at the level of organismal fitness it was shown that the hill contains a depression which makes the peak selectively inaccessible (Lunzer *et al*., 2005).

## 3. Previously noted difficulties

The idea of the adaptive landscape has met with critique from several directions (Pigliucci, 2008; Wilkins & Godfrey-Smith, 2009). Most notable is perhaps that fitness landscapes usually are visualised as 3-dimensional continuous Euclidean surfaces, while they in reality are discrete spaces with thousands of dimensions (corresponding to the number of genes considered). We can easily imagine how a population, represented by a cloud of points, moves on a landscape resembling a real mountain, but transferring this way of thinking to a space with completely different topological properties easily leads erroneous conclusions guided by our intuition of Euclidean space. Concepts such as “peaks” and “ridges” might not transfer well into the actual structure of the fitness landscape, and this might in turn lead to misconceptions about the evolutionary dynamics occurring on the landscape, such as Wright’s initial “problem” with peak-shifting.

Another issue concerns the number of dimensions (i.e. nucleotides, genes or traits) needed for a faithful description of the system. Optimally every gene would be taken into account, however that is not feasible, so a balance must be struck between the completeness and tractability of the model. With respect to traits, yet another problem arises, which relates to how traits are defined and to what degree they can be separated (Gould & Lewontin, 1979).

The landscape metaphor in most instance assumes a constant and fixed environment, which defines the fitness of the organisms. This is problematic in three ways; firstly it ignores the possibility of a changing environment, such as slow changes to the climate which might affect the adaptive value of the organisms. Viewed from this perspective the shape of the fitness landscape ought to be time-dependent, and should also depend on abiotic factors such as climate, or on changes in the ecosystem, such as the introduction of new species which might compete or predate on the organisms in question (Bak & Sneppen, 1993).

On a smaller scale it fails to capture the fact that the fitness of an individual might depend on the number of other organisms which inhabit the same point in configuration space, a phenomenon known as frequency-dependent selection, which acts to maintain variability in the population, and is for example the cause of colour polymorphism observed in many prey organisms (Bond, 2007). This suggests that the adaptive landscape should rather be thought of as a soft structure which changes its shape depending on the distribution of the population.

Further, the metaphor neglects the possibility that the organisms alter their environment in a beneficial direction, known as niche construction (Odling-Smee *et al*., 2003). This is a common feature, and is exemplified by beaver dams, which not only provide necessary shelter for the beavers themselves, but also substantially change the surrounding ecosystem, which in turn affects later generations of beavers. This is obviously not an instantaneous process, and implies that the shape of the fitness landscape depends on the history of the population and how it previously occupied the configuration space (Lewontin, 1983).

Lastly, it does not take into account the effect of genetic drift, i.e. the tendency of deleterious mutations by chance being fixed in an evolving population. This effect is prevalent in small populations where stochastic effects in reproduction are enhanced. There has been a long-standing debate over the relative importance of adaptation vs. drift (Nei, December 2005), and although this has not been resolved it has been shown that genetic drift is an important factor in molecular evolution (Kimura, 1983). From this perspective the fitness landscape does not give an accurate description of the evolutionary dynamics for small population, as genetic drift can lead to movement away from peaks in the landscape.

This summarises the previously raised issues with fitness landscapes, and we will now move on to the problem of directional variation, but before doing so we must introduce a couple of mathematical concepts, which will illuminate the subject.

## 4. Metrics

The notion of distance in genotype and phenotype space lie at the very core of the landscape metaphor, and we will therefore in this section describe how they are defined and related. Mathematically, distance in a topological space *X* is defined by a metric, a function *d*: *X* × *X* → ℝ which satisfies the following conditions:

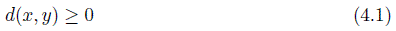

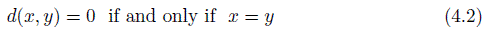

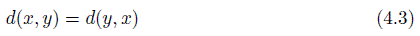

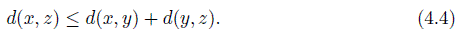

The first condition states that the distance between any two points is always positive or zero. The second states that two points are at distance zero if and only if they are identical, the third condition states that the distance is symmetric, and finally the fourth states what is known as the triangle inequality.

### (a) Metrics in genotype space

In the case of genotypes the notion of “nearness” or distance is naturally induced by the genetic operators which generate variation in the population (Shpak & Wagner, 2000). Two genotypes are considered as neighbours in the configuration space if one can be generated from the other, and vice versa, through successive generations. This idea has been formalised in the language of pretopologies (Stadler *et al*., 2002), which in a mathematical framework captures the way new variants are generated from a given set of genotypes. In order to understand this treatment we need to introduce some formalism.

For simplicity we assume a binary genome **G** consisting of 1’s and 0’s of a given length *N*, i.e. **G** = (1101010….10), which we can think of as for example RNA sequences consisting of only guanine and uracil (Stadler, 2006). Please note that before we specify any genetic operators acting on the binary genomes the collection of all genotypes is not endowed with any structure, but simply corresponds to the set of all binary sequences of length N, of which there are 2^*N*^. The simplest possible genetic operator, which is known to exists in all organisms, is a point mutation, which randomly changes one base pair (or bit in our case) of the genotype into another. This operation induces a structure on the configuration space, where two sequences are said to be adjacent if they can be reached by a single point mutation, and this structure is known as the hypercube of dimension *N*. The metric in this case is the Hamming distance, which counts the total number of positions in which two sequences differ, and is easily seen to fulfill the requirements of a metric (eq. 4).

Diploid organisms are also subject to recombination of genetic material, known as chromosomal cross-over, which is also an important genetic operator. It works by breaking up two matching chromosomes and joining them to create two novel combinations of genes on the pair of chromosomes. If we for simplicity consider recombination of our previously defined binary genomes, and restrict ourselves to homologous recombination (i.e. equal cross-over, which results in two equal sized chromosomes), then the cross-over operator can be formalised as function 𝓡: *X* × *X* → 𝓟(*X*), that takes pairs of genomes into the power set (set of all subsets) of *X*, and for a given pair of genotypes returns the set of all genomes that can be generated by cross-over from the considered pair. Although the topological structure in this case is more complicated, it can still be shown that the recombination space can be given a metric (Shpak & Wagner, 2000), which is related to the size of the recombinant set of two genotypes. The logic behind this is that the more similar two genotypes are the lower number of different recombinant genotypes can be created from them.

These genetic operators thus give rise to genotype spaces which can be endowed with a metric, but there are other genetic changes which do not have this property. One example is unequal cross-over, where the chromosomes are not properly aligned during cross-over, resulting in two chromosomes of unequal size. A mathematical model of this process was analysed by Shpak & Wagner (2000), and they conjectured that no metric exists for this configuration space. In the next section we will give more examples of genetic operators which do not generate a metric space, but before then we move on to discuss metrics in phenotype spaces, which turns out to be more problematic.

### (b) Metrics in phenotype space

When considering adaptive landscapes which are defined in terms of phenotypic traits, the corresponding phenotypes are usually ordered in terms of some representation, e.g. increasing limb length, degree of colouring etc. This structuring of the phenotype space is based solely on the characteristics of the phenotype, and takes no consideration of the underlying genetics which give rise to the phenotype. Changes in phenotype are always induced by mutations on the genetic level, and the structure of the phenotype space should therefore reflect these underlying constraints. This relation between genotype and phenotype is usually denoted the the genotype-phenotype (GP) map Φ: *X* → *Y*, which maps all possible to genotypes in *X* to corresponding phenotypes in *Y* (assuming a constant environment). This view is of course a gross over-simplification of the incredibly complex developmental process multi-cellular organisms go through, but all the same it is a highly useful conceptual tool for understanding phenotypic evolution. The full structure of the GP-map is only known in the simplest of biological systems, the folding of RNA sequences (genotypes) into shapes or secondary structures (phenotypes) (Stadler, 2006). The main features of this mapping are: (i) the GP-map is many-to-one, i.e. many genotypes map to the same phenotype, (ii) many secondary structures are rare and few are common, and (iii) sequences folding into a common secondary structure are connected into what is termed a neutral network. These properties are also believed to hold for more complicated developmental systems, and give a hint of the structure of more complex GP-maps, whose elusiveness lie in the fact that they cross many spatial and temporal scales in the developmental process from a single cell to an entire organism.

The structure of the phenotype space is thus induced by the structure of the genotype space via the GP-map, in the sense that only phenotypes whose corresponding genotypes are close in the genotype space are considered close to each other. This idea has been formalised by Stadler *et al*. (2001), allowing them to define concepts such as independence of characters and continuity in a rigorous fashion. The treatment here of phenotype spaces will be informal, but it is worth mentioning that even if the underlying genetic space is metric only particular choice of GP-maps will render the phenotype space metric.

The problem with this approach is that it requires full, or at least partial knowledge, of the GP-map of the organism in question, and this is still lacking in most cases. The success of traditional phenotypic fitness landscapes also shows that it is a feasible approach, but in order to understand why these simplified models work it is important to have a firm grip of theoretical basis of the concept.

## 5. Absence of metrics

In this section we will present evidence, on both the phenotypic and genotypic scale, that many of configuration spaces fail to incorporate a metric in the topological sense. From this follows, that in these spaces no natural notion of distance exists and consequently any analogy to Euclidean space is flawed. This however, does not imply that the metaphor of a fitness landscape in these instances is lost all together; it only means that we have to modify our notion of how fitness landscapes are defined and how this influences the evolutionary dynamics which occurs on them.

### (a) Genotype spaces

Many of the processes which generate genetic variation and novelty have an underlying nature of asymmetry to them. If a process occurs more readily in one direction than the other then there is a built-in bias which drives the system towards one state rather than the other. Further, the path taken by the system might influence future biases, so that the dynamics of the system also becomes path-dependent.

A striking example of this is whole-genome duplication which is known to have occurred in the lineages of many extant organisms such as in the ancestor of *S. cerevisiae* (Cliften *et al*., 2006). This genetic mechanism is more prevalent among plants where it is believed to be a major mechanism in speciation (Wood *et al*., 2009), and in particular among ferns (Wagner & Wagner, 1979), where some species exhibit three or more sets of chromosomes. It has also been argued that genome duplications are responsible for many major transitions in the history of life, such as the Cambrian explosion (Meyer & Schartl, 1999), and that it might be correlated with increase in organismal complexity and species diversity (Crow & Wagner, 2006). The process of genome duplication occurs through failed meiosis, producing diploid gametes, which then self-fertilise to produce a tetraploid organism. This process occurs quite readily, especially in ferns, while the reverse, i.e. the loss of half of the genetic material in one generation is virtually impossible. Naturally loss of genes does occur, for example *S. cerevisiae* has lost 88 % of the genes generated by its ancestral whole genome duplication (Kellis *et al*., 2004), but the important thing to stress is that it does not occur through a reversal of the duplication process, but that the genes are lost one at a time, in a completely different process (Krylov *et al*., 2003).

Less dramatic than the duplication of the entire genome is the change in numbers of single chromosomes. Such chromosomes that exhibit fluctuations in abundance are usually known as B-chromosomes and are prevalent in plants (Camacho *et al*., 2000), but have also been found in animals, such as the grasshopper *Eyprepocnemis plorans* (Lopez-Leon *et al*., 1992). The function of these supernumerary chromosomes is in most cases unclear, although in a few cases positive fitness effects have been observed (Jones & Rees, 1982). However, it is not their effect on fitness, or if they are fixed in the population which concerns us, but instead the underlying genetic processes which increases or decreases their number, and it has been shown that they tend to accumulate, and that their numbers in natural populations are kept down not by the removal of chromosomes, but by selection against individual carrying many supernumeraries. Again this points to an asymmetric process, where going from an organism with *n* chromosomes to one with *n* + 1 occurs at a higher rate than the reverse process.

The main mechanisms for generating genetic novelty is not the duplication of entire chromosomes, but the duplication of chromosome segments, which results in the duplication of the genes within that segment (Ohno, 1970; Zhang, 2003). This can occur by two main mechanisms; unequal cross-over, and retroposition. The former consists of a recombination event in which the chromosomes are not properly aligned, while the latter occurs when RNA of a gene is retrotranscribed into DNA, which is then inserted at a new location in the genome. When duplication of a gene has taken place the selection pressure on that gene is in most cases low due to the redundancy in functionality, although in some cases several identical copies of a gene can be advantageous. This means that the new copy can acquire mutations which can have a number of different consequences. If deleterious mutations accumulate, the gene loses its functionality and turns into a pseudogene, which is then either deleted or becomes what is usually termed “junk-DNA”. However, in some cases beneficial mutations can lead to sub-functionalisation or even neo-functionalisation of the duplicated the gene, in which case it is retained. It has been estimated that duplicate genes are formed at a rate of 1 /gene/100 MY (million years) in eukaryotes (Lynch & Conery, 2000), while the rate of gene loss is estimated to be 10 times higher (Lynch & Conery, 2003). This means that the process of gene duplication is an asymmetric process for generating variation, and in a sense even irreversible, since returning to the exact same genotype which existed prior to the duplication event is highly unlikely, as genes are lost by point-mutations which lead to a gradual decay, and not by sudden excisions.

A further example of a genetic process which violates the condition of symmetry is the lateral (or horizontal) transfer (LGT) of genes among bacteria (Ochman *et al*., 2000). This process was detected through the rapid spread of multi-drug resistance among diverse species of bacteria in the 1950s (Davies, 1996), and has since been recognised as the main mechanisms of bacterial innovation. The amount of genetic material acquired by LGT in *E. coli* has been estimated to be 16 % (Ochman *et al*., 2000) of all genetic material, and the rate of gene acquisition in the *Streptococcus* genus has been estimated to be larger than the per nucleotide substitution rate, which highlights its significance. There is also evidence that more insertion than deletions occur, at least in more recent bacterial species (Hao & Golding, 2004), which again implies an asymmetric structure of the genotype space.

Not even point mutations, which previously were taken as a sound example of non-directional genetic change, are completely symmetric. It has been shown that the rate of mutation differ significantly depending on the location in the DNA, being influenced by the base composition of the genes (Wolfe *et al*., 1989). This means that certain genes are more prone to mutation, and that variation therefore occurs more readily along certain axes in the genetic landscape. There is also a bias in the rate of transversion between different bases, such that the GC → AT mutation rate is higher than the AT → GC mutation rate (Ellegren *et al*, 2003).

From this brief review it becomes clear that directional variation is present on all scales where genetic change occurs, from the drastic duplication of entire genomes to point mutations, the smallest unit of genetic variation.

The above processes of variation are all genetic changes which induce a structure on the genotype space that is more or less directional, meaning that mutating from genotype A to B might occur much more readily than going from B to A. Formally A and B are still neighbours in the genotype space considered, but the rate at which B organisms are generated from A organisms is not the same as from B to A. To put things clearly the mutation rate *µ_A,B_* from A to B is, in the cases we have discussed, not equal to *µ_B,A_*. The simplified genotype spaces discussed in sec. b do not have this property, as all point mutations are equally like, i.e. *µ_ij_* = *µ_ji_* for all *i* and *j*, and the same is true for all possible recombination events. The only reasonable way to capture this directionality is to say that the distance between A and B is not the same as B to A, and this relation contradicts the third requirement of a metric (eq. 4.3).

If we now suppose that we, on top of the imagined genotype space, construct a fitness landscape, which to each genotype assigns an elevation reflecting its reproductive success, then this landscape indeed will have some strange properties. The main problem lies in the fact that it cannot be visualised in an unambiguous manner, as the distances in the landscape depend from where they are perceived. There is, so to speak, no objective way to view the topography. From this follows that the idea of the population performing a hill-climbing motion towards local peaks in the landscape is a misconception, as each individual might perceive a different landscape. This idea will be developed further and formalised in section 6, but first we will tackle the problems encountered in phenotype space.

### (b) Phenotype spaces

Depending on the structure of the GP-map the phenotype space might inherit some of the directional variation discussed in the previous chapter. Some genetic changes might be neutral in which case they will not affect the phenotype, but in the cases where genotypic changes are not neutral the directionality of the genotype space is carried over into phenotype space.

However, the are other cases where asymmetries exist in phenotype space and these are related to the structure of the GP-map itself, and do not depend on the processes generating variation *per se*. For clarity, consider an RNA-sequence which can fold into two distinct shapes depending on the base pair sequence. It is a well known fact that sequence-to-shape mappings are many to one, i.e. many different nucleotide sequences fold into the same shape, and following this we can safely assume that the number of sequences which fold into one of the shapes of our hypothetical secondary shape is larger than the other.

When this system is viewed at the genotype level together with the knowledge of the folding map, the evolutionary dynamics would be perfectly explicable. However, when viewed only at the shape or phenotype level, without the knowledge of the underlying sequences, it would seem as if one of the shapes was preferred over the other, as if there was a bias (or one sequence was selected for). Assuming complete ly random *genetic* mutations, the *phenotypic* changes would seem biased, a phenomenon usually labelled directional mutation (Maynard Smith *et al*., 1985), and leads to “ascent of the abundant” a term coined by Cowperthwaite *et al*. (2008) in relation to RNA sequence evolution.

The example of RNA evolution was intentionally chosen at the lowest possible level of biological organisation, but already here we observe a directionality in the process of phenotypic change. Moving up the temporal and spatial scales to organisms and to the kind of individual-level traits often used in fitness landscapes the degree of complication is not likely to lessen.

A different form of directionality relates to developmental constraints found among multi-cellular organisms. The amount of possible phenotypic variation is often constrained by the developmental pathway of the organism (Maynard Smith *et al*., 1985) or by pleoitropic gene effects (Arnold, 1994). This suggests that some hypothetical phenotypes, which from a purely visual inspection would be said to lie close to a given wild-type, in fact are very unlikely or even impossible to reach. Let us give an example of this, which relates to phalanx reduction among frogs (eutetrapods) and salamanders (urodeles), and is caused by the developmental differences between the two orders. Experimental work has show that, by reducing the number of cells available for digit development, the first digit to be lost is the last one to develop (Wiens & Hoverman, 2008), and because eutetrapods and urodeles have different developmental pathways (the order in which the digits appear) different patterns in digit loss are observed. The loss of digits itself is driven by natural selection, but the pattern by which it occurs is dictated by the developmental constraints. The point here is that a phenotype might seem close from a morphological perspective, but is very far from a genetic and ontogenic point of view. This also highlight how developmental pathways might phenotypic change might constrain the dynamics to a subspace of the phenotype space whose dimensionality depends on the number of independent traits or characters (Wagner & Stadler, 2003).

## 6. A dynamical approach

The genetic mechanisms discussed in the previous sections, which generate variation in a population, form one corner stone in evolution, and are the means by which any type of movement occurs on the fitness landscape. The other mechanisms are natural selection, which guarantees that the general direction of movement is up the gradients in the landscape, and drift which adds a component of random movement to the population. An approach which incorporates all these processes is to view evolution as a random walk on a fitness landscape (with asymmetric) jump probabilities (Sella & Hirsh, 2005).

Assuming that mutation rate is small we can consider a situation where the population is monomorphic at any given time and look at the probability of fixation of a mutant genotype *j* in the resident wild-type population *i*. Using an approximation of the Fisher-Wright process, the probability of fixation is given by,

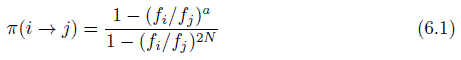

where *f_i_* is the fitness of genotype *i*, *N* is the effective population size, and *a* = 2 in haploid populations and *a* = 1 in diploid. This expression only incorporates the effects of selection and drift, and the complete expression of the rate of transition is given by

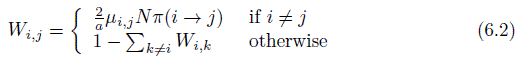

where *µ_ij_* is the mutation rate from genotype *i* to *j*, which is only non-zero if the genotypes are neighbours in the space of genotypes considered. For a given initial state this equation defines a Markov chain which describes the probability *P_i_*(*t*) of finding the population fixed for genotype *i* at time *t*, and evolves according to

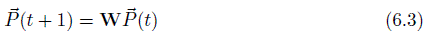

where 
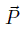
 is a probability vector and **W** is the matrix of transition rates. Under the condition the Markov chain is irreducible a unique stationary distribution 
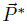
 exists which satisfies the condition 
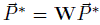
. The *i*th component of this vector corresponds to the probability of finding the population in a state where genotype *i* is fixed. One option is to take this distribution 
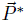
 to be a representation of the fitness landscape, which incorporates the effect of mutation, selection and drift.

If the dimensionality of the fitness landscape is large then visualising this vector becomes problematic. A solution to this problem was proposed by McCandlish (2011) who suggested that the eigenvectors of the transition matrix **W** could be used as a basis for visualising the structure of the genotype space. That procedure places genotype *i* at x-coordinate *u*_2_(*i*) and y-coordinate *u*_3_(*i*), where *u*_2,3_ are the second and third largest eigenvectors of **W**. The mutational structure is introduced by connecting two genotypes with a line if they can be reached by a single mutation. Lastly the fitness of each genotype can be visualised by mapping it to the size of the node representing each genotype.

## 7. Conclusion

We have in this paper pointed out the lack of a proper metric in many genotypic and phenotypic configurations spaces, and thus an absence of a notion of distance in the corresponding fitness landscapes. This is rooted in the lack of symmetry in several mechanisms generating genetic variation, such as whole genome duplication and lateral gene transfer. The asymmetry in phenotype space stems from these underlying genetic processes, but also from properties of the GP-map and developmental constraints. These facts taken together shows that ordinary fitness landscapes might be poor representations of the true evolutionary process, in particular if mutational asymmetries are abundant.

This situation can however be remedied. Instead of viewing fitness landscapes simply as being shaped by fitness, one should also take into account how mutation rates and the effects of genetic drift impact evolution.

## Footnotes

† We will use the term ‘fitness landscape’, however the two terms have been used interchangeably in the literature.

